# A single closed head injury in mice induces chronic, progressive white matter atrophy and increased phospho-tau expressing oligodendrocytes

**DOI:** 10.1101/2022.05.19.492705

**Authors:** David F. Havlicek, Rachel Furhang, Elena Nikulina, Bayle Smith-Salzberg, Siobhán Lawless, Sasha A. Sevarin, Sevara Mallaboeva, Fizza Nayab, Alan C. Seifert, John F. Crary, Peter J. Bergold

## Abstract

Traumatic brain injury (TBI) acutely damages the brain; this injury can evolve into chronic neurodegeneration. While much is known about the chronic effects arising from multiple mild TBIs, far less is known about the long-term effects of a single moderate to severe TBI. We found that a single moderate closed head injury to mice induces diffuse axonal injury within 1-day post-injury (DPI). At 14 DPI, injured animals have atrophy of ipsilesional cortex, thalamus, and corpus callosum, with bilateral atrophy of the dorsal fornix. Atrophy of the ipsilesional corpus callosum is accompanied by decreased fractional anisotropy and increased mean and radial diffusivity that remains unchanged between 14 and 180 DPI. Injured animals increased density of phospho-tau immunoreactive (pTau^+^) cells in the ipsilesional cortex and thalamus, and bilaterally in corpus callosum. Between 14 and 180 DPI, atrophy occurs in the ipsilesional ventral fornix, contralesional corpus callosum, and bilateral internal capsule. Diffusion tensor MRI parameters remain unchanged in white matter regions with delayed atrophy. Between 14 and 180 DPI, pTau^+^ cell density increases bilaterally in corpus callosum, but decreases in cortex and thalamus. The location of pTau^+^ cells within the ipsilesional corpus callosum changes between 14 and 180 DPI; density of all cells increases including pTau^+^ or pTau^-^ cells. Greater than 90% of the pTau^+^ cells are in the oligodendrocyte lineage in both gray and white matter. Density of thioflavin-S^+^ cells in thalamus increases by 180 DPI. These data suggest a single closed head impact produces multiple forms of chronic neurodegeneration. Gray and white matter regions proximal to the impact site undergo rapid atrophy. More distal white matter regions undergo chronic, progressive white matter atrophy with an increasing density of oligodendrocytes containing pTau. These data suggest that the chronic neurodegeneration arising from a single moderate CHI differs greatly from the chronic traumatic encephalopathy produced by multiple mild head injuries.

**Highlights:** Gray and white matter atrophy begins within 14 days after a single closed head injury

White matter atrophy progresses between 14 and 180 days post injury with minimal changes in diffusion tensor MRI parameters.

CHI increases the density of oligodendrocytes with perinuclear accumulation of phosphorylated tau

Thioflavin-S^+^ cells increase in thalamus at 180 days post injury

## Introduction

Acute damage following a TBI may develop into a chronic disease (Graham and Sharp, 2019; Johnson et al., 2017; Pattinson and Gill, 2018; Walker and Tesco, 2013; Wilson et al., 2017). Forty percent of TBI patients suffer from progressive deficits in coordination, information processing, and executive function that either plateau or slowly worsen years after a moderate to severe TBI (Vaishnavi et al., 2009). The high prevalence of chronic degenerative outcomes suggests a single TBI can readily develop into a progressive neurodegenerative disorder (Graham and Sharp, 2019; Tsai et al., 2021). TBI can be roughly divided into acute, subacute and chronic phases (Algattas and Huang, 2013; Harris et al., 2019; Tsai et al., 2021). An acute phase ends days after injury, an subacute phase lasts weeks after injury, and a chronic phase lasts months to years. As yet, these phases are not strictly defined by any biochemical, radiological or metabolic outcomes (Algattas and Huang, 2013; Osier et al., 2015; Tsai et al., 2021). Diffuse axonal injury occurring soon after injury is a major driver of the chronic effects of TBI (Graham et al., 2020; Harris et al., 2019).

The extent of diffuse axonal injury observed soon after a single moderate to severe TBI predicts chronic neurodegeneration (Graham et al., 2020; Tsai et al., 2021). Moderate to severe TBI also induces chronic and progressive brain atrophy, with the cortex, thalamus and corpus callosum most commonly effected (Harris et al., 2019; Moen et al., 2014). Longitudinal MRI studies of clinical TBI reveals ongoing white matter atrophy with decreased fractional anisotropy and increased mean diffusivity (Harris et al., 2019; Moen et al., 2014). The corpus callosum is particularly prone to TBI damage due to the orientation and length of its axonal tracts (Harris et al., 2019; Kumar et al., 2009; Kumar et al., 2010; Moen et al., 2014; Rutgers et al., 2008)

Repetitive mild or subconcussive TBIs can slowly develop into a chronic traumatic encephalopathy (CTE) characterized by brain atrophy with hyperphosphorylated tau (pTau) aggregates in neurons and astrocytes (Kriegel et al., 2018; McKee et al., 2013). While studies of CTE have improved our understanding of how mild repetitive TBIs induce chronic neurodegenerative disease, few studies have examined the chronic outcomes following a single moderate to severe TBI. When reviewed in 2015, less than 11% of preclinical TBI studies examined outcomes more than 6 months post-injury (Osier et al., 2015).

Clinical TBI can produce chronic and progressive gray and white matter atrophy (Bigler, 2021; Harris et al., 2019). Diffusion tensor-MRI (DTI) has detected persistant and progressive alterations of white matter microstructure after clinical TBI (Hutchinson et al., 2018). In preclinical models, progressive atrophy and neural loss peristed for one year followng fluid percussion injury (Smith et al., 1997). Some rodent TBI models show only transient changes in DTI that return to sham levels after 30 DPI (Soni et al., 2020); others have shown chronic degeneration particularly of white matter (Armstrong et al., 2016; Mac Donald et al., 2007; Mohamed et al., 2021; Soni et al., 2020). In the controlled cortical impact (CCI) model of TBI, increased gray matter atrophy ocurred close to the impact site (Mohamed et al., 2021). CCI also acutely altered fractional anistoropy (FA), mean diffusivity (MD), axial difusivity (AD) and radial diffusivity (RD) in multiple white matter regions. Progressive changes in these diffusion parameters at 60 DPI suggests ongoing neurodegeneration (Mohamed et al., 2021). Pathological tau aggregates are seen in white matter injured by a single moderate to severe TBI, suggesting pTau accumulation in different locations than CTE (Gorgoraptis et al., 2019). Limited studies suggest that pTau accumulates in oligodendrocytes after a single TBI (Irving et al., 1996; Smith et al., 2003). In contrast, CTE is characterized by pTau accumulation in neurons and astrocytes (McKee et al., 2013). The presence of oligodendrocytic pTau aggregates rules out a single diagnosis of CTE (McKee et al., 2016). These data suggest that a single moderate to severe TBI produces neurodegeneration that differs from CTE.

Cytoplasmic pTau aggregates likely damage cells by impairing transport, and the integrity of cytoskeleton and mitochondria (Kametani and Hasegawa, 2018; Wang and Mandelkow, 2016). pTau aggregates are seen in multiple neurodegenerative diseases including CTE, Alzheimer’s disease, progressive supranuclear palsy, corticobasal degeneration, agyrophilic grain disease, Pick’s disease, primary age related tauopathy, and frontotemporal dementia with parkinsonism-17 (Chung et al., 2021; Clavaguera et al., 2013; Crary et al., 2014; Ferrer et al., 2014; Goedert et al., 2017; Kahlson and Colodner, 2015; LoPresti, 2018; Narasimhan et al., 2017). Depending upon the tauopathy, pTau aggregates are expressed by neurons, astrocytes or oligodendrocytes (Chung et al., 2021; Clavaguera et al., 2013; Ferrer et al., 2014; Kahlson and Colodner, 2015; LoPresti, 2018; Narasimhan et al., 2017). pTau accumulates in oligodendrocytes in corticobasal degeneration, progressive supranuclear palsy, argyrophilic grain disease, and frontotemporal dementia with parkinsonism-17 (LoPresti, 2018; Richter-Landsberg, 2016). Expression of mutant pathogenic human tau in murine oligodendrocytes damaged axons and myelin, and increased motor deficits (Higuchi et al., 2005). Intracerebral injection of pTau from the oligodendrocytes of patients with corticobasal degeneration or progressive supranuclear palsy induced pTau aggregation in the oligodendrocytes of recipient mice (Narasimhan et al., 2020). These pTau expressing oligodendrocytes could transmit pTau to neighboring oligodendrocytes (Narasimhan et al., 2020). pTau aggregates that spread to neighboring cells contain a pathogenic β-pleated sheets that bind the fluorescent dye, thioflavin S (DeVos et al., 2017; Nizynski et al., 2017; Santa-María et al., 2006).

Few histologic studies have evaluated the regional and cellular distribution of pTau after a single TBI in humans (Irving et al., 1996; Johnson et al., 2012; Smith et al., 2003; Zanier et al., 2018). Irving et al. utilized co-labeling studies to demonstrate an increase in Tau expression in oligodendrocytes acutely after a single TBI or stroke in humans (Irving et al., 1996). Smith et al. reported increased Tau expression in oligodendrocytes less than 1 month after a single TBI in humans, yet did not report an increase in Tau expression in neurons relative to non-TBI controls (Smith et al., 2003). Tau aggregates were reported in neurons and glia after a single TBI in humans, yet the cellular identity of the glial tau expressers was not reported (Johnson et al., 2012; Zanier et al., 2018). pTau^+^ neurons and glia were reported in cerebral cortex, hippocampus, thalamus and corpus callosum 3 months and 12 months after a single CCI to mice, but the glial cell type was not characterized (Zanier et al., 2018). pTau^+^ oligodendrocytes were seen after a single blast injury, and caspase-3-cleaved tau was observed in the corpus callosum after a single CCI (Du et al., 2016; Glushakova et al., 2018). Several animal studies have also reported an increase in Tau^+^ oligodendrocytes after experimental stroke (Irving et al., 1997; Sun et al., 2002; Valeriani et al., 2000).

This study uses a closed head injury (CHI) model to study chronic effects of TBI (Grin’kina et al., 2016). During CHI, a piston directly strikes the head of the mouse, producing large acceleration-deceleration movements while compressing the brain through the skull. This produces a transient loss of righting reflex, a rapid induction of hemorrhage, and contusion proximal to the impact site (Grin’kina et al., 2016; Whitney et al., 2021). Gray and white matter injury are induced both proximal and distal to the impact site (Grin’kina et al., 2016; Sangobowale et al., 2018; Whitney et al., 2021). CHI produces contusion and hemorrhage with transient loss of consciousness suggesting that it models moderate TBI. In this study, MRI and histology were used to study the subacute and chronic changes following CHI.

## Methods

### Mouse closed head injury (CHI) model of TBI

Male C57BL/6J mice (16 weeks old, 28-30g) received either sham closed head injury (sham CHI) or closed head injury (CHI), as described previously (Sangobowale et al., 2018). Upon arrival to SUNY-Downstate, mice received a unique identification number. All subsequent determinations were done by an observer blinded to the treatment received by the mice. Briefly, deep anesthesia was induced (3% isoflurane in oxygen (2.0 L/min)) and maintained (2% isoflurane in oxygen (2.0 L/min)) until immediately after impact. The shaved mouse head was placed on a modified Kopf stereotaxic apparatus with its bed and ear bar holders covered with 0.5 inches of polyurethane foam. The right hemisphere was impacted with a 4.0 mm diameter impactor tip placed 3 mm lateral from the midline and 3 mm rostral from the anterior end of the ears. Using an electromagnetic impactor, a single 6.4 m/s impact was delivered that compressed the skull to a depth of 3 mm with dwell time of 1 second. The head moved freely during and after impact (Grin’kina et al., 2016). After CHI, mice were placed on a heating pad until the animal resumed an upright position and then returned to their home cages. Sham CHI used the same procedure without an impact. All animal studies were conducted at State University of New York (SUNY) Downstate Health Sciences University and approved by the Institutional Animal Care and Use Committee of SUNY Downstate (Animal welfare assurance A3260-01; Protocol 19-10571). All studies were performed in accordance with the NIH *Guide for the Care and Use of Laboratory Animals*.

### Histology

At 14 or 180 days post-injury (DPI), mice were deeply anesthetized with isoflurane (3%) in oxygen (2 L/min) and perfused transcardially with paraformaldehyde (4% (w/v)) in phosphate buffered saline (PBS). The brain was collected and treated with 4% paraformaldehyde for 24 hours at 4°C. Following transfer to PBS, brains were stored at 4°C for up to 1 month prior to processing and paraffin embedding. Parasagittal sections (5μm) are prepared 1.6 ± 0.6 mm from the midline. Chromogenic immunohistochemistry was performed on a Ventana Benchmark XT according to manufacturer’s directions. Sections were processed using CC1 antigen retrieval buffer (Tris/Borate/ EDTA buffer, pH 8.0–8.5, Roche Diagnostics, Basel Switzerland). Table 1 contains details of the antibodies used in this study. Primary antibodies were diluted in antibody dilution buffer (ABD24, Roche Diagnostics) and detected with the UltraView or OptiView secondary detection kits (Roche Diagnostics). Antibody specificity was confirmed by staining slides containing brain sections from the CHI or sham CHI groups as well as a section from a tauopathy patient and an aged control patient. All the pTau antibodies tested (AT8, S214, RZ3, PHF1, and PG5) showed cellular immunoreactivity in the tauopathy, but not the control sections (not shown). Brain sections from the CHI and sham CHI groups showed minimal staining with RZ3 and PG5 (not shown). Slides were counterstained with hematoxylin. Immunoblots of thalamic protein showed that AT8, PHF1 and S214 recognized proteins having the reported molecular weight of pTau (Supplementary Figure 1). Cells positive for S214, PHF1, AT8, Olig2 or Thioflavin-S were assessed in cortex (all values are mm from Bregma;, AP -1.0 - -0.54; DV, 0.5 - 1.2; ML, 0.8 - 1.8), thalamus (AP, -0.6 - -3.0; DV, 2.0 -3.6; ML, 0.8 – 1.8) and corpus callosum (AP, -1.0 - -0.65; DV, 1.0 - 1.2; ML, 0.8 - 1.8). Regional densities of S214, PHF1, and AT8 in corpus callosum were assessed in parasagittal sections located in four 600µm regions located anterior from the impact site (AP, 1.4 – 1.0). Bielschowsky’s silver stain was performed on 8μm parasagittal sections located 1.6mm ± 1.2mm from midline as described previously. An Aperio CS2 slide scanner scanned and digitized slides stained using chromogenic immunohistochemistry. Slides were incubated with Thioflavin S (0.05**%**, MedChemExpress) in the dark for 5 minutes at room temperature washed with 70% ethanol for 2 minutes, and distilled water twice for 2 minutes. Fluorescent stained slides were scanned and digitized using a Zeiss Axio Observer LSM 800 microscope.

**Table 1.**
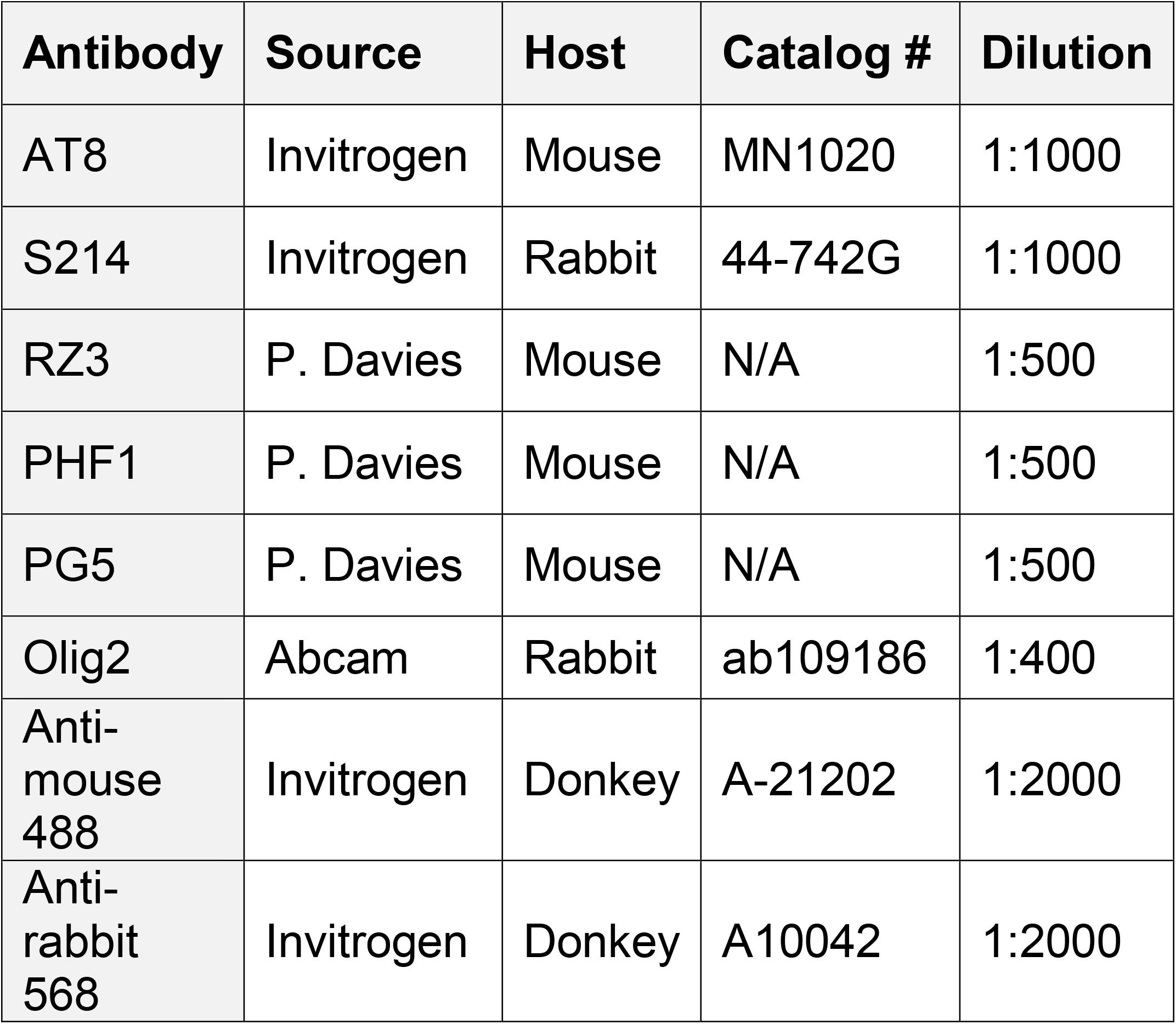
Antibodies used in this study

QuPath quantified axonal bulbs, pTau^+^ cells or thioflavin-S in digitally scanned slides (Version: 0.3.0) (Bankhead et al., 2017). For each analysis, regions of interest were created manually. Scripts used to detect positive cells are in a GitHub repository located at this link: https://github.com/Bergold-Lab/QuPath-Positive-Cell-Identification.

### Magnetic resonance imaging acquisition and analysis

Magnetic resonance imaging (MRI) was performed on a 9.4 T Bruker Avance III 400 micro-MRI system using a 20-mm quadrature radiofrequency coil. Formalin fixed mouse brains were immersed in PBS in 20-mm glass tubes, agitated under vacuum to dislodge residual air bubbles, and imaged using T2-weighted and diffusion-weighted sequences. Anatomical T2-weighted imaging was performed using a 3D rapidly-refocused echo (RARE) sequence with the following parameters: 125 μm isotropic resolution, field of view = 20mm*20mm*12mm, repetition time = 2 s, effective echo time = 17.5 ms, RARE factor = 4, averages = 2. Diffusion weighted images (DWI) were acquired using a diffusion-weighted spin-echo sequence with the following parameters: 172 µm isotropic resolution, field of view =20mm*20mm*12mm, repetition time = 5 s, echo time = 18.05 ms, averages = 10, 6 diffusion encoding directions at b = 1000 s/mm2 and 1 b = 0 image. Images were co-registered and parcellated using AIDAmri. Volumes of the corpus callosum (0 to 2.7 DV (from Bregma (in mm)); cortex (0 to -3.0 AP and 1.4-1.7 L); thalamus (2.36-4.28 DV); internal capsule (2.68-4.88 DV), dorsal fornix (1.60-2.04 DV), and ventral fornix (2.80-5.36 DV))(Paxinos and Franklin, 2019) were quantified using Mango 4.1 image viewer software. Diffusion-weighted data were pre-processed using the FMRIB Software Library (FSL)’s eddy current correction tool, ‘eddy’. Diffusion tensor imaging (DTI) analysis was performed using FSL ‘dtifit’. Ipsilesional and contralesional corpus callosum volumes between 0 to 2.7mm from Bregma were determined using the FSL package’s image viewer, FSLeyes, on a virtual workstation (VMware Workstation 16). Diffusion microstructural parameters were then calculated in regions of interest defined using FSLeyes.

### Statistics

Two-tailed unpaired T-tests were used to analyze all pairwise comparisons. Regional pTau^+^ cell density differences in ipsilesional corpus callosum were analyzed using three-way multivariate analysis. Effects of injury, time and their interaction were evaluated using two-way ANOVA. If two-way ANOVA or three-way multivariable analysis showed a significant injury effect, one-way ANOVA and Tukey’s multiple comparisons post-hoc test evaluated for injury effects at 14 and 180 DPI. The full statistics of the two-way ANOVA are provided in Tables 2, 3 and 4, only the one-way ANOVA and post-hoc results are in the text. Ipsilesional thalamus volume and thioflavin S positive cells were first analyzed by Pearson correlation followed by a linear regression. Statistical significance was set at 0.05. Exact probability values are provided unless the value is less than 0.001. All data is reported as mean ± SEM.

**Table 2.**
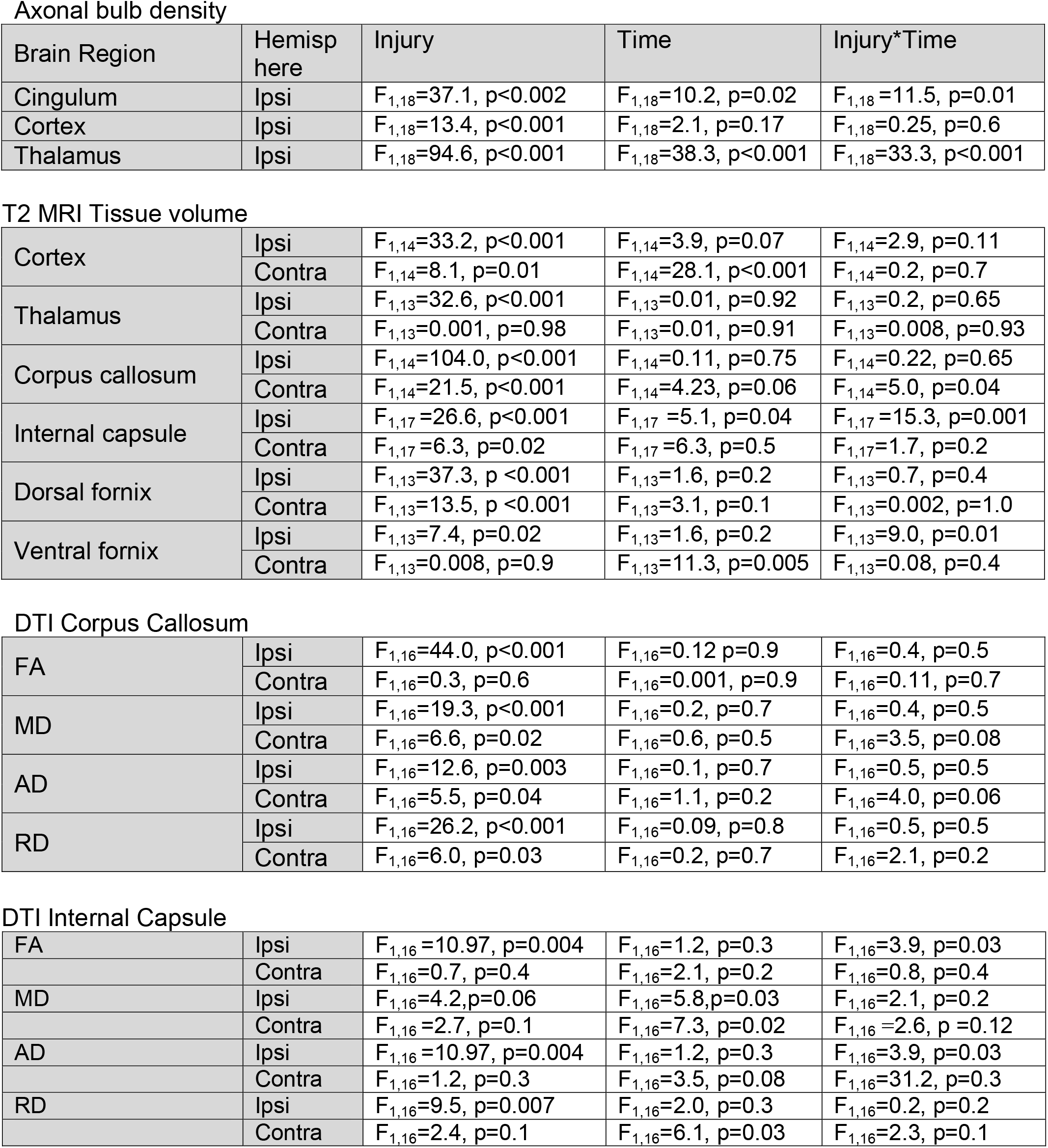
ANOVA Table of Axonal Retraction Bulb Density and MRI measurements. Ipsi, ipsilesional; Contra, contralesional

Axonal bulb density
| Brain Region | Hemisp here | Injury | Time | Injury*Time |
| --- | --- | --- | --- | --- |
| Cingulum | Ipsi | $F_{1,18}=37.1, p<0.002$ | $F_{1,18}=10.2, p=0.02$ | $F_{1,18}=11.5, p=0.01$ |
| Cortex | Ipsi | $F_{1,18}=13.4, p<0.001$ | $F_{1,18}=2.1, p=0.17$ | $F_{1,18}=0.25, p=0.6$ |
| Thalamus | Ipsi | $F_{1,18}=94.6, p<0.001$ | $F_{1,18}=38.3, p<0.001$ | $F_{1,18}=33.3, p<0.001$ |
| T2 MRI Tissue volume |  |  |  |  |
| Cortex | Ipsi | $F_{1,14}=33.2, p<0.001$ | $F_{1,14}=3.9, p=0.07$ | $F_{1,14}=2.9, p=0.11$ |
| | Contra | $F_{1,14}=8.1, p=0.01$ | $F_{1,14}=28.1, p<0.001$ | $F_{1,14}=0.2, p=0.7$ |
| Thalamus | Ipsi | $F_{1,13}=32.6, p<0.001$ | $F_{1,13}=0.01, p=0.92$ | $F_{1,13}=0.2, p=0.65$ |
| | Contra | $F_{1,13}=0.001, p=0.98$ | $F_{1,13}=0.01, p=0.91$ | $F_{1,13}=0.008, p=0.93$ |
| Corpus callosum | Ipsi | $F_{1,14}=104.0, p<0.001$ | $F_{1,14}=0.11, p=0.75$ | $F_{1,14}=0.22, p=0.65$ |
| | Contra | $F_{1,14}=21.5, p<0.001$ | $F_{1,14}=4.23, p=0.06$ | $F_{1,14}=5.0, p=0.04$ |
| Internal capsule | Ipsi | $F_{1,17}=26.6, p<0.001$ | $F_{1,17}=5.1, p=0.04$ | $F_{1,17}=15.3, p=0.001$ |
| | Contra | $F_{1,17}=6.3, p=0.02$ | $F_{1,17}=6.3, p=0.5$ | $F_{1,17}=1.7, p=0.2$ |
| Dorsal fornix | Ipsi | $F_{1,13}=37.3, p<0.001$ | $F_{1,13}=1.6, p=0.2$ | $F_{1,13}=0.7, p=0.4$ |
| | Contra | $F_{1,13}=13.5, p<0.001$ | $F_{1,13}=3.1, p=0.1$ | $F_{1,13}=0.002, p=1.0$ |
| Ventral fornix | Ipsi | $F_{1,13}=7.4, p=0.02$ | $F_{1,13}=1.6, p=0.2$ | $F_{1,13}=9.0, p=0.01$ |
| | Contra | $F_{1,13}=0.008, p=0.9$ | $F_{1,13}=11.3, p=0.005$ | $F_{1,13}=0.08, p=0.4$ |
| DTI Corpus Callosum |  |  |  |  |
| FA | Ipsi | $F_{1,16}=44.0, p<0.001$ | $F_{1,16}=0.12, p=0.9$ | $F_{1,16}=0.4, p=0.5$ |
| | Contra | $F_{1,16}=0.3, p=0.6$ | $F_{1,16}=0.001, p=0.9$ | $F_{1,16}=0.11, p=0.7$ |
| MD | Ipsi | $F_{1,16}=19.3, p<0.001$ | $F_{1,16}=0.2, p=0.7$ | $F_{1,16}=0.4, p=0.5$ |
| | Contra | $F_{1,16}=6.6, p=0.02$ | $F_{1,16}=0.6, p=0.5$ | $F_{1,16}=3.5, p=0.08$ |
| AD | Ipsi | $F_{1,16}=12.6, p=0.003$ | $F_{1,16}=0.1, p=0.7$ | $F_{1,16}=0.5, p=0.5$ |
| | Contra | $F_{1,16}=5.5, p=0.04$ | $F_{1,16}=1.1, p=0.2$ | $F_{1,16}=4.0, p=0.06$ |
| RD | Ipsi | $F_{1,16}=26.2, p<0.001$ | $F_{1,16}=0.09, p=0.8$ | $F_{1,16}=0.5, p=0.5$ |
| | Contra | $F_{1,16}=6.0, p=0.03$ | $F_{1,16}=0.2, p=0.7$ | $F_{1,16}=2.1, p=0.2$ |
| DTI Internal Capsule |  |  |  |  |
| FA | Ipsi | $F_{1,16}=10.97, p=0.004$ | $F_{1,16}=1.2, p=0.3$ | $F_{1,16}=3.9, p=0.03$ |
| | Contra | $F_{1,16}=0.7, p=0.4$ | $F_{1,16}=2.1, p=0.2$ | $F_{1,16}=0.8, p=0.4$ |
| MD | Ipsi | $F_{1,16}=4.2, p=0.06$ | $F_{1,16}=5.8, p=0.03$ | $F_{1,16}=2.1, p=0.2$ |
| | Contra | $F_{1,16}=2.7, p=0.1$ | $F_{1,16}=7.3, p=0.02$ | $F_{1,16}=2.6, p=0.12$ |
| AD | Ipsi | $F_{1,16}=10.97, p=0.004$ | $F_{1,16}=1.2, p=0.3$ | $F_{1,16}=3.9, p=0.03$ |
| | Contra | $F_{1,16}=1.2, p=0.3$ | $F_{1,16}=3.5, p=0.08$ | $F_{1,16}=31.2, p=0.3$ |
| RD | Ipsi | $F_{1,16}=9.5, p=0.007$ | $F_{1,16}=2.0, p=0.3$ | $F_{1,16}=0.2, p=0.2$ |
| | Contra | $F_{1,16}=2.4, p=0.1$ | $F_{1,16}=6.1, p=0.03$ | $F_{1,16}=2.3, p=0.1$ |

**Table 3.**
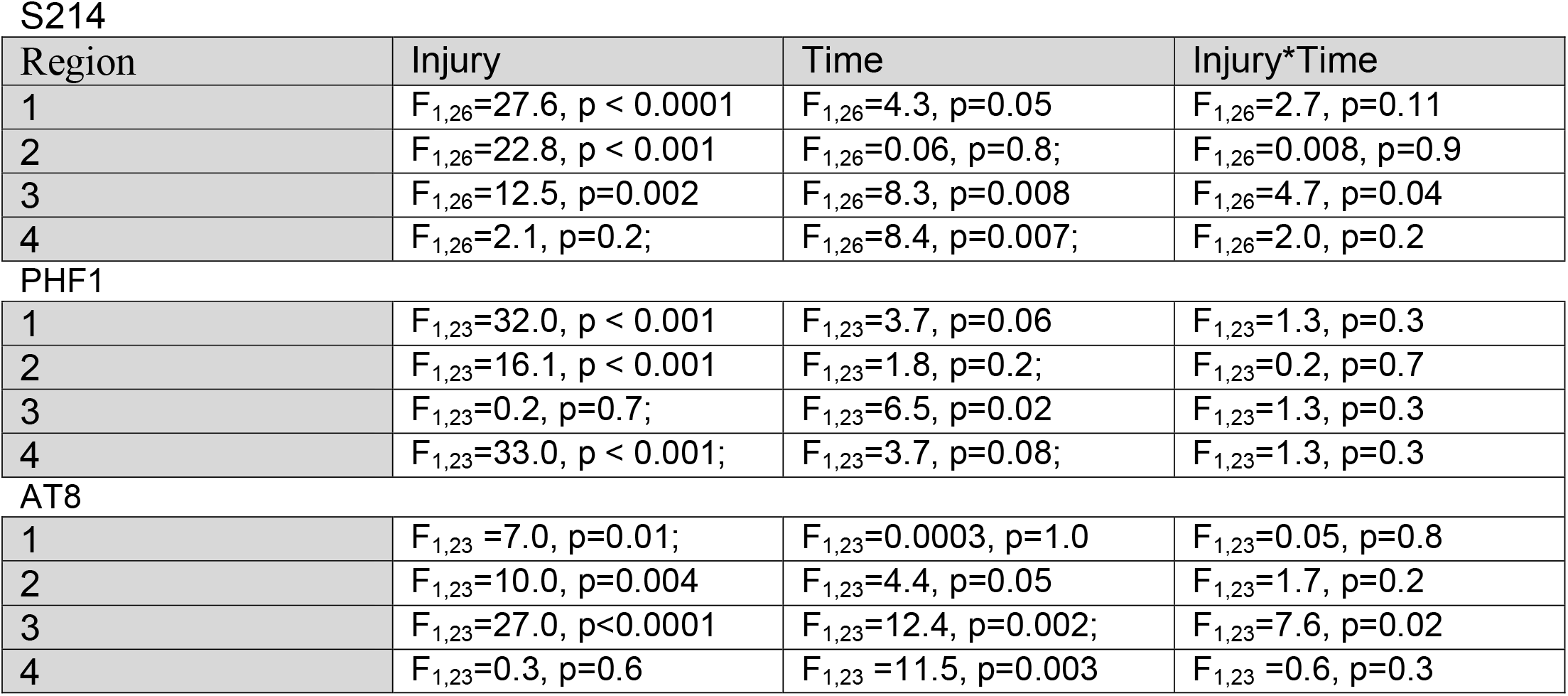
ANOVA table of regional changes in pTau^+^ cell density in ipsilesional corpus callosum

| Region | Injury | Time | Injury*Time |
| --- | --- | --- | --- |
| 1 | $F_{1,26}=27.6, p < 0.0001$ | $F_{1,26}=4.3, p=0.05$ | $F_{1,26}=2.7, p=0.11$ |
| 2 | $F_{1,26}=22.8, p < 0.001$ | $F_{1,26}=0.06, p=0.8;$ | $F_{1,26}=0.008, p=0.9$ |
| 3 | $F_{1,26}=12.5, p=0.002$ | $F_{1,26}=8.3, p=0.008$ | $F_{1,26}=4.7, p=0.04$ |
| 4 | $F_{1,26}=2.1, p=0.2;$ | $F_{1,26}=8.4, p=0.007;$ | $F_{1,26}=2.0, p=0.2$ |
| PHF1 |  |  |  |
| 1 | $F_{1,23}=32.0, p < 0.001$ | $F_{1,23}=3.7, p=0.06$ | $F_{1,23}=1.3, p=0.3$ |
| 2 | $F_{1,23}=16.1, p < 0.001$ | $F_{1,23}=1.8, p=0.2;$ | $F_{1,23}=0.2, p=0.7$ |
| 3 | $F_{1,23}=0.2, p=0.7;$ | $F_{1,23}=6.5, p=0.02$ | $F_{1,23}=1.3, p=0.3$ |
| 4 | $F_{1,23}=33.0, p < 0.001;$ | $F_{1,23}=3.7, p=0.08;$ | $F_{1,23}=1.3, p=0.3$ |
| AT8 |  |  |  |
| 1 | $F_{1,23} = 7.0, p=0.01;$ | $F_{1,23}=0.0003, p=1.0$ | $F_{1,23}=0.05, p=0.8$ |
| 2 | $F_{1,23}=10.0, p=0.004$ | $F_{1,23}=4.4, p=0.05$ | $F_{1,23}=1.7, p=0.2$ |
| 3 | $F_{1,23}=27.0, p<0.0001$ | $F_{1,23}=12.4, p=0.002;$ | $F_{1,23}=7.6, p=0.02$ |
| 4 | $F_{1,23}=0.3, p=0.6$ | $F_{1,23} = 11.5, p=0.003$ | $F_{1,23} = 0.6, p=0.3$ |

**Table 4.**
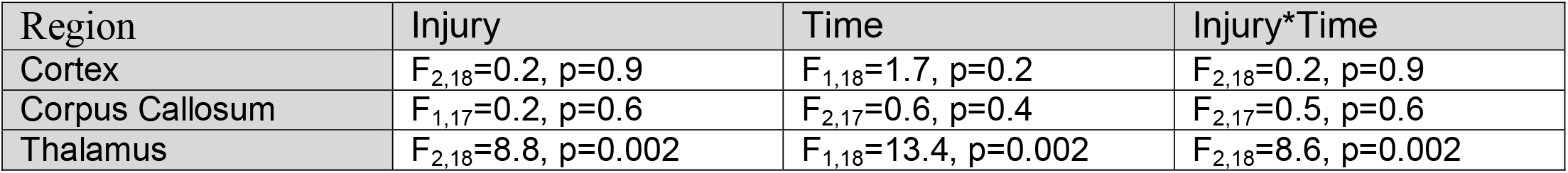
ANOVA table of changes in thioflavin-S^+^ cell density

| Region | Injury | Time | Injury*Time |
| --- | --- | --- | --- |
| Cortex | $F_{2,18}=0.2, p=0.9$ | $F_{1,18}=1.7, p=0.2$ | $F_{2,18}=0.2, p=0.9$ |
| Corpus Callosum | $F_{1,17}=0.2, p=0.6$ | $F_{2,17}=0.6, p=0.4$ | $F_{2,17}=0.5, p=0.6$ |
| Thalamus | $F_{2,18}=8.8, p=0.002$ | $F_{1,18}=13.4, p=0.002$ | $F_{2,18}=8.6, p=0.002$ |

## Results

### CHI rapidly induces axonal damage in both gray and white matter

Rapid and delayed axonal damage occurs in both clinical TBI and in animal models of TBI (Armstrong et al., 2016; Greer et al., 2011; Johnson et al., 2017). Axotomy produces axonal retraction bulbs that are visualized using Bielschowsky’s silver stain (Thomas et al., 2018; van de Looij et al., 2012). Silver-stained axonal bulbs were assessed in thalamus, cingulum, and cortex of sham and injured mice at 1 and 14 DPI (Figure 1). In the cingulum, axonal bulb density had a significant effect of injury, time and an interaction of injury and time (injury*time). At both 1 and 14 DPI, axonal bulb density in the cingulum of injured mice was significantly higher than sham injured mice (1 DPI, p< 0.001; 14 DPI, p=0.05). The density of axonal bulbs in injured mice decreased significantly between 1 and 14 DPI (p=0.008).

**Figure 1.**
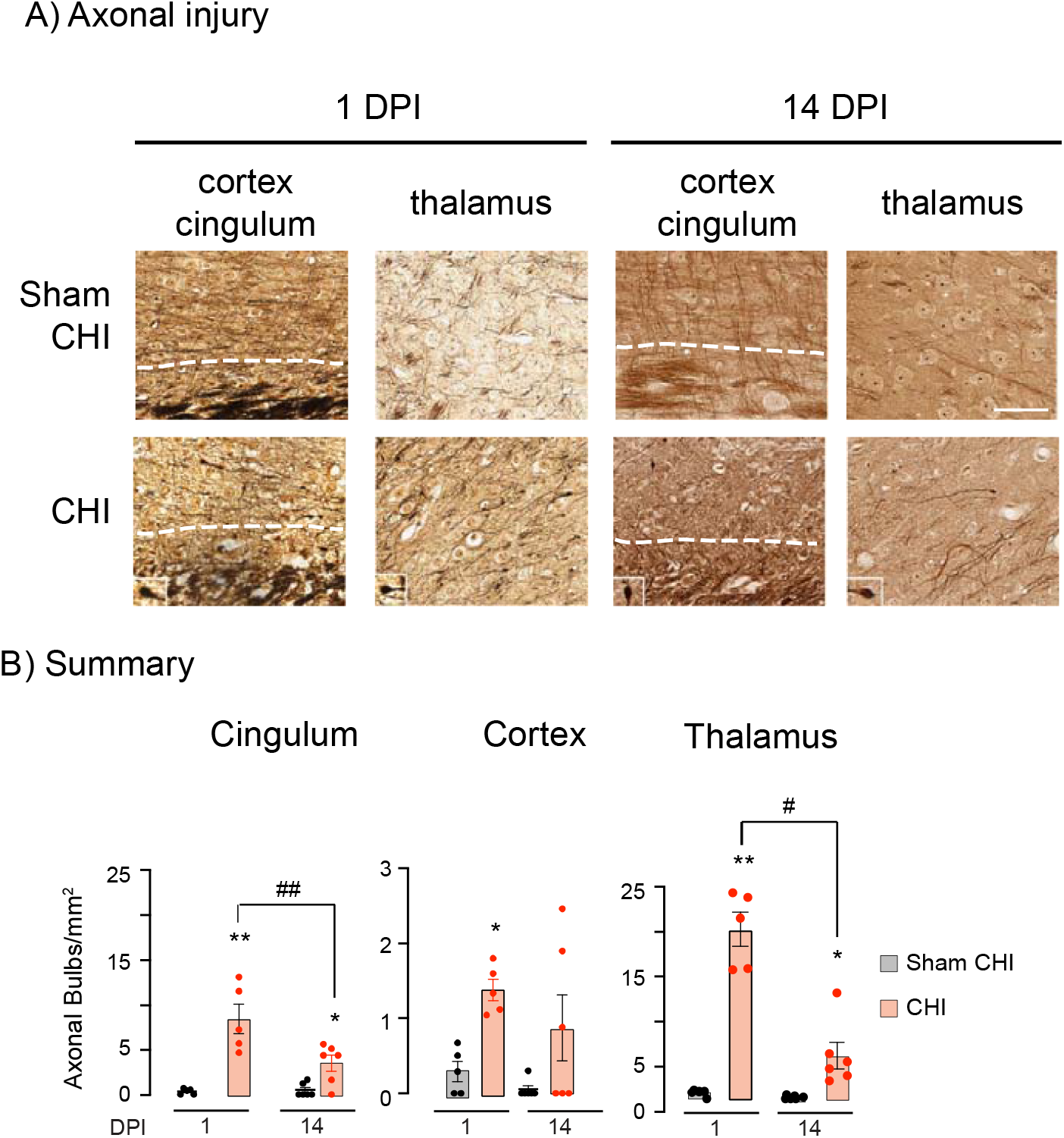
Axonal injury following CHI. Panel A, Bielschowsky’s silver stained axons from the cortex, cingulum and thalamus of sham-CHI or CHI mice. A dotted line delineates the cortex from the underlying cingulum and corpus callosum. An insert on the lower left shows representative axonal bulbs from each section. Scale Bar for all panels 10µm. Panel B, Summary of axon bulb density. At 1 DPI, the cingulum, cortex and thalamus of CHI mice had a higher density of axonal bulbs than sham-CHI mice; at 14 DPI, cingulum and thalamus of CHI mice had a higher density of axonal bulbs than sham-CHI mice (*p<0.05; **p<0.001). At 1DPI, the cingulum and thalamus of CHI mice had more axonal bulbs than at 14 DPI (*p<0.05; **p<0.01).

Cortical axonal bulb density showed a significant effect of injury. At 1 DPI, injured mice had higher axonal bulb density in the cortex than sham injured mice (p=0.03). At 14 DPI, axonal bulb density in the cortex of injured mice trended to be higher than in sham treated mice (p=0.06).

In the thalamus, axonal bulb density had significant effects of injury, time and injury*time. Axonal bulb density in the thalamus of injured mice was significantly higher than in sham injured mice at both 1 and 14 DPI (1 DPI, p< 0.001; 14 DPI, p=0.02). Axonal bulb density in injured mice significantly decreased between 1 and 14 DPI (p=0.05). These data suggest that axonal injury occurred rapidly in the cingulum, cortex, and thalamus. Axonal injury persisted in cingulum and thalamus at 14 DPI.

### CHI induces rapid atrophy of gray and white matter followed by delayed, progressive atrophy of white matter

The volumes of the cortex, thalamus, corpus callosum, internal capsule and fornix were assessed at 14 and 180 DPI by *ex vivo* T2-weighted MRI (Figure 2). Ipsilesional cortical volume had a significant effect of injury. At both 14 and 180 DPI, injured mice had a smaller ipsilesional cortex than sham inured mice (F_3,14_=12.8, p < 0.001, 14 DPI, p <0.001; 180 DPI, p=0.04). Contralesional cortical volume had significant effects of injury and time. Sham CHI mice developed a contralesional cortical atrophy between 14 and 180 DPI (p=0.006). CHI mice also showed age-dependent atrophy (F_3,14_=12.04, p < 0.001; p=0.02). Cortical volume was smaller in CHI mice than in sham CHI mice at 180 DPI (p>0.001). These data suggest injured mice had an early ipsilesional cortical atrophy that remained unchanged after 14 DPI. In contrast, contralesional cortex had a delayed, age- and injury-dependent atrophy beginning later than 14 DPI.

**Figure 2.**
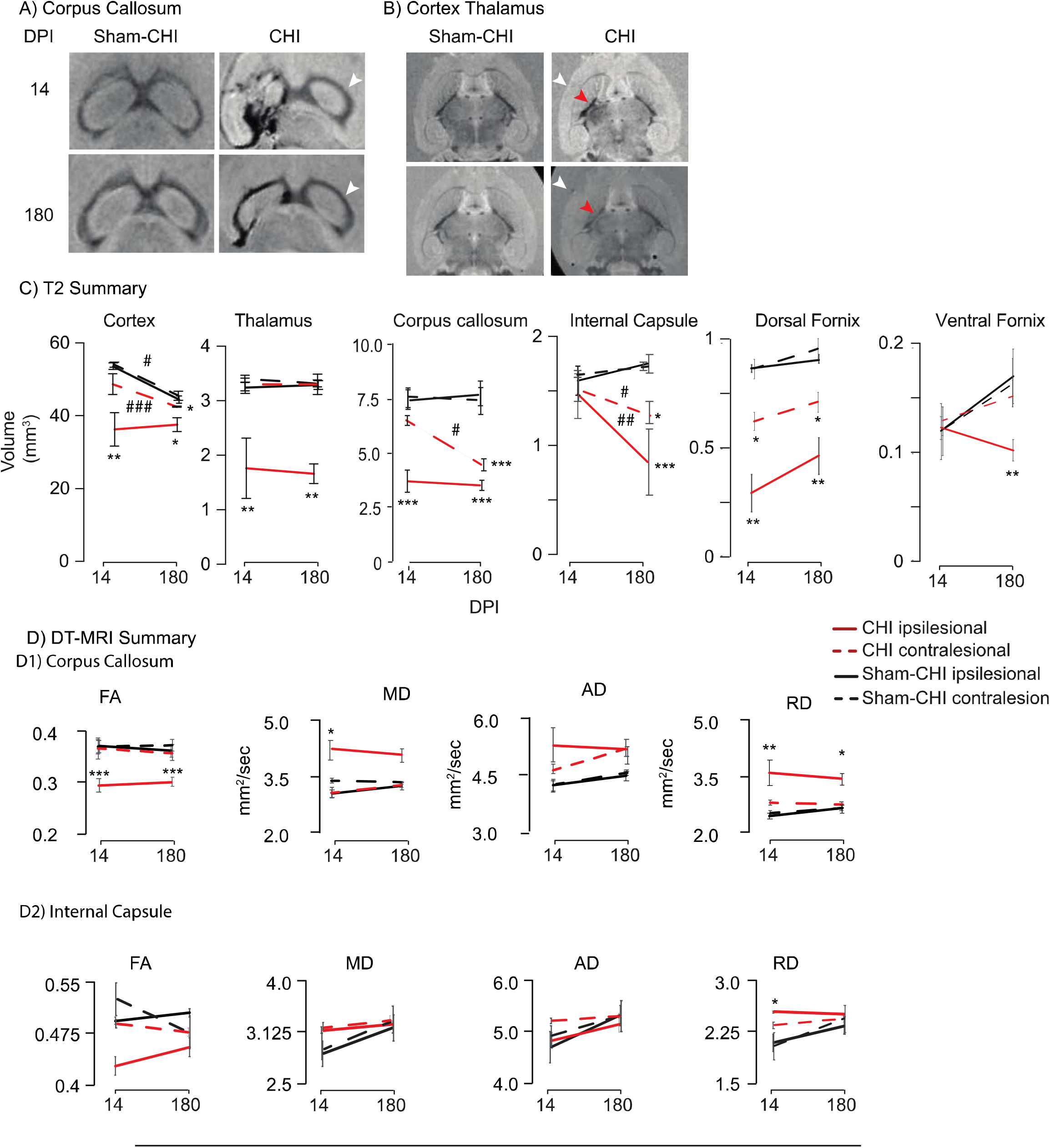
Rapid and delayed atrophy after a single CHI. Panel A, Representative horizontal *ex vivo* T2-weighted MRI images of corpus callosum in mice at 14 or 180 days after receiving either sham-CHI or CHI. An arrowhead points to an area of contralesional of corpus callosum showing atrophy at 180 DPI. Panel B, Representative horizontal *ex vivo* T2-weighted MRI images of cortex and thalamus in mice at 14 or 180 days after receiving either sham-CHI or CHI. Arrowheads shows atrophy of ipsilesional cortex (white) or internal capsule (red). Panel C, Summary of ipsilesional and contralesional volume changes in multiple gray and white matter regions. CHI induced delayed atrophy between 14 and 180 DPI in contralesional cortex and corpus callosum, bilaterally in internal capsule and ipsilesionally in ventral fornix. Panel D, Summary of *ex vivo* DT-MRI. D1, Corpus callosum, at 14 DPI, FA, MD, and RD of the ipsilesional corpus callosum differed between CHI and sham-CHI mice; at 180 DPI, FA and RD differed between CHI and sham-CHI mice. At 14 DPI, MD of the contralesional corpus callosum differed between CHI and sham-CHI mice (*p < 0.05, **p < 0.01; ***p < 0.001). D2, Internal Capsule, at 14 DPI, CHI mice had a higher RD than sham-CHI mice (*p=0.05). Asterisks indicate significant difference from sham-CHI (*p < 0.05, **p <0.01, ***p < 0.001) and hashtags indicate significant differences between 14 and 180 DPI (^#^p < 0.05, ^##^p < 0.01, ^###^p < 0.001).

Ipsilesional thalamic volume had a significant effect of injury. Ipsilesional thalamus of CHI mice was smaller than sham CHI mice at both 14 DPI and 180 DPI (F_3,13_=11.06, p < 0.001; 14DPI, p=0.02; 180 DPI, p=0.003). Contralesional thalamic volume had no significant effects. These data suggest that CHI produced ipsilesional thalamic atrophy that was unchanged between 14 and 180 DPI.

Ipsilesional corpus callosum volume had a significant injury effect. Ipsilesional corpus callosum of injured mice was smaller than sham CHI mice at both 14 and 180 DPI (F_3,14_=35.58, p < 0.001, 14 DPI, p < 0.001; 180DPI, p < 0.001). Contralesional corpus callosum volume had significant injury and injury*time effects that trended toward significance with time. At 180DPI, contralesional corpus callosum of injured mice was smaller than sham CHI mice (F_3,14_=11.71, p < 0.001, 180DPI, p < 0.001). Contralesional corpus callosum atrophy increased in CHI mice between 14 and 180 DPI (p=0.04). These data suggest an early atrophy of ipsilesional corpus callosum and a delayed and progressive atrophy in contralesional corpus callosum.

Ipsilesional internal capsule volume had significant effects of injury, time and injury*time. Ipsilesional nternal capsule of the CHI mice was smaller than sham CHI mice at both 14 and 180 DPI (F_3,17_ =20.4, p<0.001; 14 DPI, p=0.002, 180 DPI p < 0.001). Contralesional internal capsule volume had a significant injury effect (F_3,17_ =3.6, p=0.04). Contralesional internal capsule volume of CHI mice at 180 DPI was smaller than sham-CHI mice (p=0.03). Contralesional internal capsule volume of CHI mice also decreased between 14 and 180 DPI (p=0.03). These data suggest delayed, progressive bilateral atrophy of internal capsule.

Ipsilesional dorsal fornix volume had a significant effect of injury. At both 14 and 180 DPI, ipsilesional dorsal fornix of the CHI group was significantly smaller than the sham CHI group (F_3,13_=9.3, p=0.001; 14 DPI, p=0.003; 180 DPI, p=0.005). Contralesional dorsal fornix volume had a significant effect of injury. At both 14 and 180 DPI, contralesional dorsal fornix of the CHI group was smaller than the sham CHI group (F_3,13_=13.5, p <0.001; 14 DPI p=0.04; 180 DPI, p=0.01). Ipsilesional ventral fornix volume had significant effects of injury and injury*time. At 180 DPI, ipsilesional ventral fornix was smaller in the CHI group than the sham CHI group (F_3,13_=7.3, p=0.004; 180 DPI, p=0.003). Contralesional ventral fornix volume was unchanged.

These data suggest that CHI induced rapid atrophy of the ipsilesional cortex, corpus callosum, and thalamus, with bilateral atrophy of the dorsal fornix. This rapid atrophy was followed by a delayed progressive atrophy of the contralesional cortex and corpus callosum, ipsilesional ventral fornix, and bilateral internal capsule (Figure 2C).

### CHI induces changes in white matter integrity

DTI assessed microstructural changes in corpus callosum and internal capsule (Figure 2D). Ipsilesional corpus callosum fractional anisotropy (FA) had a significant injury effect. At both 14 DPI and 180 DPI, ipsilesional corpus callosum FA of injured mice was lower than sham CHI mice (F_3,16_=14.9, p < 0.001; 14 DPI, p=0.001; 180 DPI, p=0.001). Contralesional corpus callosum FA was unchanged.

Mean diffusivity (MD) in the ipsilesional corpus callosum had a significant injury effect. At 14 DPI, but not 180 DPI, CHI increased ipsilesional corpus callosum MD (F_3,16_=6.6, p=0.004, 14 DPI, p=0.02;). Contralesional corpus callosum MD had a significant injury effect. At 14 DPI, CHI decreased contralesional corpus callosum MD (F_3,16_=3.1, p=0.06, 14 DPI p=0.05).

Ipsilesional corpus callosum axial diffusivity (AD) had a significant injury effect. At both 14 DPI and 180 DPI, ipsilesional corpus callosum AD of injured mice strongly trended to increase at 14 DPI that did not increase further at 180 DPI (F_3,16_=3.7, p=0.03;). Contralesional corpus callosum AD also had a significant injury effect. At 14 DPI, contralesional corpus callosum AD in the CHI group trended to increase (F_3,16_=2.9, p=0.07, 14 DPI, p=0.07).

Ipsilesional corpus callosum radial diffusivity (RD) had a significant injury effect. Ipsilesional corpus callosum RD increased in injured mice at 14DPI and 180 DPI (F_3,16_=8.9, p=0.001, 14 DPI, p=0.008; 180 DPI, p=0.02). Contralesional corpus callosum RD had a significant injury effect (F_3,16_=2.4, p=0.05), but no pairwise comparisons were significant.

Ipsilesional internal capsule FA has a significant effect of injury. No pairwise comparisons were significant despite a significant injury effect (F_3,16_=3.9, p=0.03). Contralesional internal capsule FA had no significant effects. Ipsilesional internal capsule MD trended toward an effect of injury (p=0.06) with a significant effect of time (p=0.03), but not injury*time with no significant pairwise comparisons. Contralesional internal capsule MD only had a significant time effect. Ipsilesional internal capsule AD only had significant effect of injury. Contralesional AD was unchanged. Ipsilesional RD has a significant effect of injury. At 14 DPI, injured mice had a higher RD than sham injured mice (F_3,16_=3.9, p=0.03, p=0.05). Contralesional RD only had a significant time effect.

### CHI induces regional increases in pTau immunoreactivity

Clinical studies suggest that pTau levels increase in areas of white matter injury after a single moderate to severe TBI (Gorgoraptis et al., 2019). Therefore, pTau expression after a single CHI was assessed using the pTau antibodies S214, PHF1 and AT8, which recognize unique tau phosphorylation sites (Goedert et al., 1995; Otvos et al., 1994; Yu et al., 2009) S214^+^, PHF1^+^ and AT8^+^ cell density was assessed in ipsilesional and contralesional cortex, corpus callosum and thalamus of sham CHI and CHI mice at 14 and 180 DPI (Figure 3).

**Figure 3.**
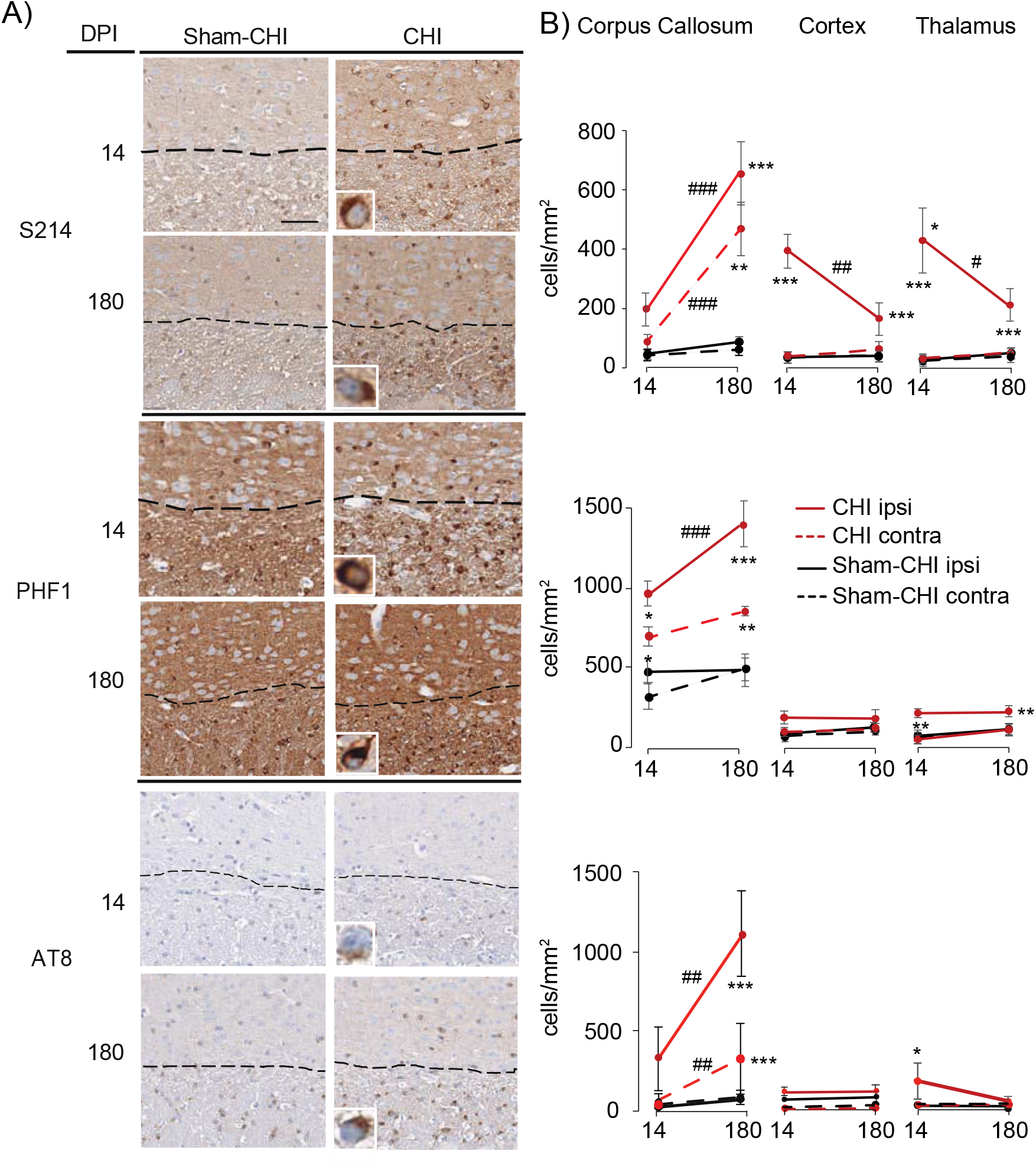
CHI increased pTau^+^ cell density in cortex, corpus callosum, and thalamus. Panel A, Representative images of the cortex and corpus callosum of sham-CHI or CHI mice stained with S214, PHF1 or AT8 anti-pTau antibodies. Scale bar 50µm. The insert of each panel shows a high magnification image of perinuclear pTau^+^ immunoreactivity. Panel B, Summary of changes in pTau^+^ cell density at 14 and 180 DPI. Asterisks indicate significant difference from sham-CHI (*p < 0.05, **p <0.01, ***p < 0.001) and hashtags indicate significant differences between 14 and 180 DPI (^#^p < 0.05, ^##^p < 0.01, ^##^p < 0.001).

S214^+^ cell density in ipsilesional corpus callosum had significant effects of injury, time and injury*time. CHI increased S214^+^ cell density at 180DPI (F_3,20_=15.8, p<0.001; 180DPI, p<0.001). S214^+^ cell density also increased between 14 and 180 DPI (p<0.001). S214^+^ cell density in contralesional corpus callosum had significant effects of injury, time and injury*time. At 180DPI, CHI significantly increased S214^+^ cell density (F_3,20_=12.6, p<0.001, p=0.003).

Between 14 and 180 DPI, S214^+^ cell density also increased (p<0.001). S214^+^ cell density in ipsilesional cortex had significant effects of injury, time and injury*time. CHI increased cortical S214^+^ cell density at 14 and 180DPI (F_3,20_=13.5, p<0.001, 14 DPI, p<0.001; 180 DPI, p<0.001). CHI mice decreased ipsilesional cortical S214^+^ cell density between 14 and 180 DPI (p=0.006). S214^+^ cell density in the contralesional cortex was unchanged. S214^+^ cell density in ipsilesional thalamus had a significant effect of injury. CHI increased S214^+^ cell density in thalamus compared to sham at 14 DPI, but not 180DPI (F_3,20_=9.7, p<0.001;14 DPI, p=0.001). S214^+^ cell density in the ipsilateral thalamus then decreased between 14 and 180DPI (p=0.05). S214^+^ cell density did not change in the contralesional thalamus. These data suggest that the ipsilesional cortex and thalamus increased S214^+^ cell density at 14 DPI that subsequently decreased by 180 DPI. These data suggest gray and white matter have different patterns of S214^+^ cell density changes.

PHF1^+^ cell density in ipsilesional corpus callosum had a significant injury effect. At 14 and 180 DPI, PHF1^+^ cell density of the CHI group was higher than the sham CHI group (F_3.14_=17.5, p < 0.001; 14 DPI, p=0.04; 180DPI, p < 0.001). CHI increased PHF1^+^cell density between 14 and 180 DPI (p < 0.001). PHF1^+^ cell density in the contralesional corpus callosum only showed an injury effect. CHI increased PHF1^+^ cell density in the contralateral corpus callosum at both 14 and 180 DPI (F_3.14_=13.1, p < 0.001; 14DPI, p=0.04; 180DPI, p =0.004).

PHF1^+^ cell density in ipsilesional and contralesional cortex had significant effects of injury but with no significant pairwise comparisons. PHF1^+^ cell density in ipsilesional thalamus had a significant injury effect. At both 14 and 180 DPI, CHI increased PHF1^+^cell density in ipsilesional thalamus (F_3,14_=8.6, p=0.002; 14 DPI, p=0.008; 180 DPI, p=0.03). PHF1^+^ cell density in the contralesional thalamus had a significant time effect with no significant pairwise comparisons.

AT8^+^ cell density in ipsilesional corpus callosum had significant effects of injury, time and injury*time. At 180 DPI, AT8^+^ cell density in the CHI group was greater than the sham CHI group (F_3,23_=17.4, p < 0.001; 180DPI, p < 0.001). CHI increased AT8^+^ cell density between 14 and 180 DPI (p=0.003). Contralesional corpus callosum AT8^+^ cell density had significant effects of injury, time, and injury*time. At 180 DPI, AT8^+^ cell density was greater in the CHI group than the sham CHI group (F_3,23_=9.9, p < 0.001; 180DPI, p < 0.001). AT8^+^ cell density in the contralateral corpus callosum increased between 14 and 180DPI (p=0.003).

AT8^+^ cell density was unchanged bilaterally in cortex. Ipsilesional thalamus had a significant injury effect. At 14 DPI, AT8+ cell density in the CHI group was greater than the sham CHI group (p=0.01). AT8+ cell density in contralesional thalamus was unchanged. Increased S214^+^, PHF1^+^ and AT8^+^ cell density in thalamus at 14 DPI were not accompanied by alterations in total thalamic pTau protein levels (Supplementary Figure 1). These data suggest a delayed increase in AT8^+^ cell density in ipsilesional corpus callosum. AT8^+^ cell density in thalamus had a modest increase followed by a delayed decrease.

### CHI produces regional differences in pTau cell density in corpus callosum

CHI increased atrophy and pTau^+^ cell density in corpus callosum (figures 2 and 3). Increased pTau^+^ cell density could occur uniformly in corpus collosum or could show regional differences (Figure 4A).

**Figure 4.**
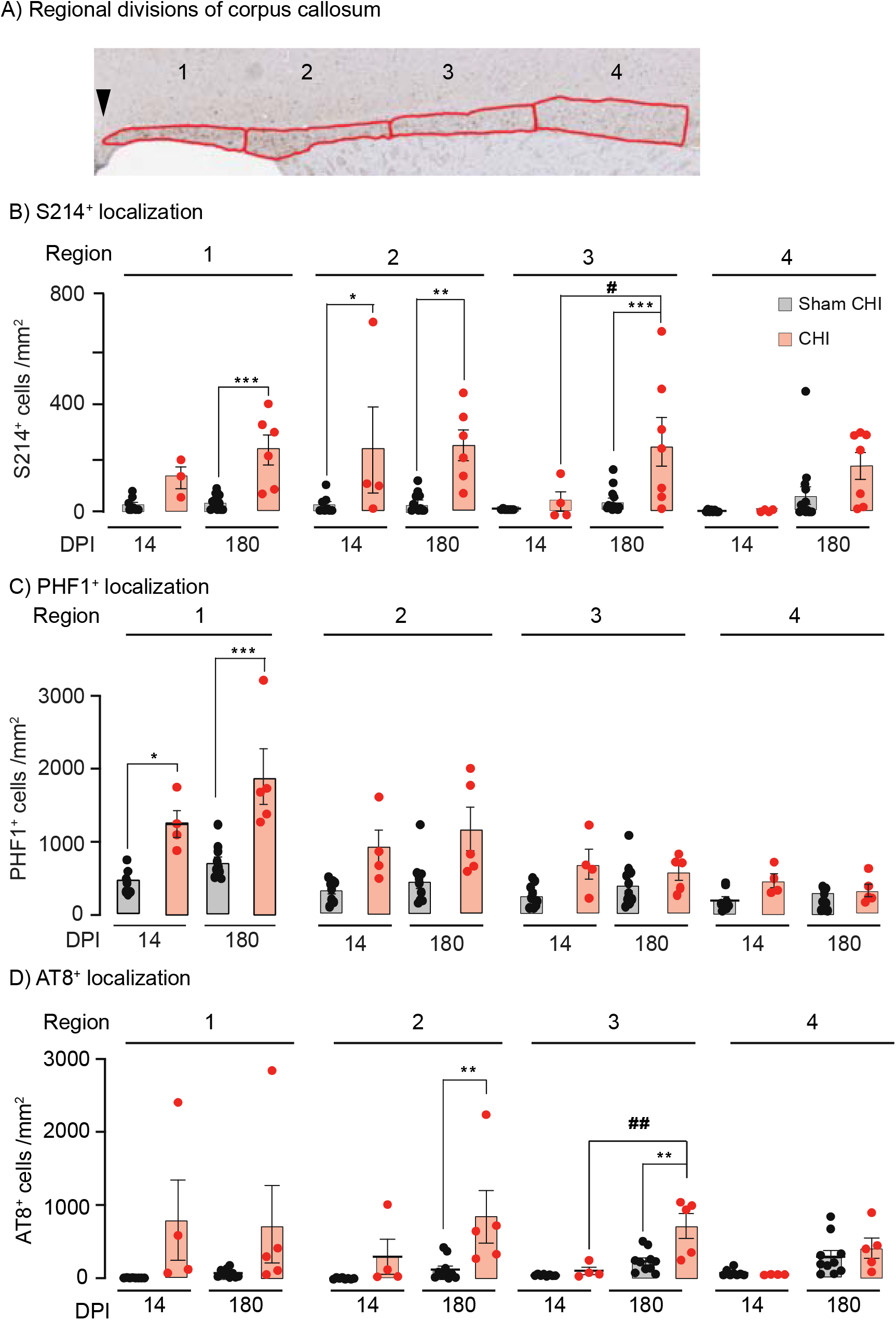
Regional changes in pTau^+^ cell density. Panel A, AT8 staining of parasagittal sections containing corpus callosum showing the impact site location (arrowhead) and the locations of the four 600µm regions located anterior to the impact site. Panel B, S214^+^ cell density significantly increased in regions 1, 2, and 3. In region 3 S214^+^ cell density increased between 14 and 180 DPI. Panel C, In region 1, PHF1^+^ cell density increased at both 14 and 180 DPI. Panel D, AT8^+^ cell density in regions 2 and 3, which had not increased at 14 DPI, significantly increased between 14 and 180 DPI. *p < 0.05, **p <

S214^+^ cell density trended strongly toward a significant interaction of region*time*injury (F_1,100_=3.47, p=0.07). S214^+^ cell density in region 1 had significant effects of injury and time, but not injury*time. CHI increased S214^+^ cell density in region 1 at 180 DPI (p<0.001). S214^+^ cell density in region 2 had a significant effect of injury, but not time or injury*time. CHI increased S214^+^ cell density at 14 (p=0.03) and 180 DPI (p=0.003). S214^+^ cell density in region 3 had significant effects of injury, time and injury*time with CHI increasing S214^+^ cell density at 180 DPI (p <0.001). S214^+^ cell density also increased between 14 and 180 DPI (p=0.02). Region 4 only had a significant effect of time. These data suggest increased S214^+^ density in region 3 between 14 and 180 DPI.

PHF1^+^ cell density trended strongly toward a significant interaction of region*time*injury (F_1,100_=3.64, p=0.06). PHF1^+^ cell density in region 1 had a significant effect of injury with a significance trend with time. CHI increased PHF1^+^ cell density at 14 (p=0.04) and 180 DPI (p < 0.0001). PHF1^+^ cell density had significant injury effects in regions 2 and 4 and a significant time effect in region 3. No significant pairwise comparisons were seen in regions 2-4. These data suggest that CHI increased PHF1^+^ cell density within region 1 of the corpus callosum at 14 and 180 DPI.

For AT8, multivariable analysis indicated a significant interaction between region*time*injury (F_1,100_=4.02, p=0.05). AT8^+^ cell density in region 1 only had a significant effect of injury, but with no significant pairwise comparisons. AT8^+^ cell density in region 2 had significant effects of time and injury. CHI increased AT8^+^ cell density at 180 DPI (p =0.006). AT8^+^ cell density in region 3 had significant effects of time, injury and injury*time. CHI increased AT8^+^ cell density at 180 DPI (p < 0.001). AT8^+^ cell density in in CHI mice was higher at 180 DPI than 14 DPI (p=0.004). AT8^+^ cell density in region 4 had a significant effect of time, but not of injury. These data suggest that regions 2 and 3 had an injury and time-dependent increase in AT8^+^ cell density.

At 180 DPI, CHI significantly increased AT8^+^ cell density in corpus callosum region 3 while total corpus callosum volume decreased (Figure 4D). To test if this increase is specific to AT8^+^ cells, total cell density was also analyzed in region 3. CHI increased the density of total cells ((in cells/mm^2^) Sham CHI 2134 ± 92.8; CHI 3629 ± 504.2; t_8_=2.9, p=0.02; These data suggest that increased AT8^+^ cell density at 180 DPI reflects an overall cell density increase.

### Oligodendrocytes aggregate pTau following CHI

The presence of perinuclear pTau aggregates in corpus callosum suggests glial pTau expression (Figure 5). Brain sections from the CHI group were co-labeled with PHF1 and Olig2, a nuclear protein expressed in cells of the oligodendrocyte lineage (Figure 5) (Rowitch et al., 2002). At 14 DPI, a large majority of PHF1^+^ cells co-labeled with Olig2^+^ in the thalamus (97.2±2.4%, n=727 cells, n=4 mice), cortex (94.4±4.5%, n=568 cells, n=4 mice) and corpus callosum (95.8 ± 3.4%, n=426 cells, n=4 mice). A similarly high percentage of co-labelling was seen at 180 DPI in thalamus (92.9 ± 21.5%, n=943 cells, n=4 mice), cortex (91.5±0.4%, n=530 cells, n=4 mice) and corpus callosum (94.0±1.7%, n=379 cells, n=4 mice). These data suggest that cells in the oligodendrocyte lineage accumulate pTau after a single CHI.

**Figure 5.**
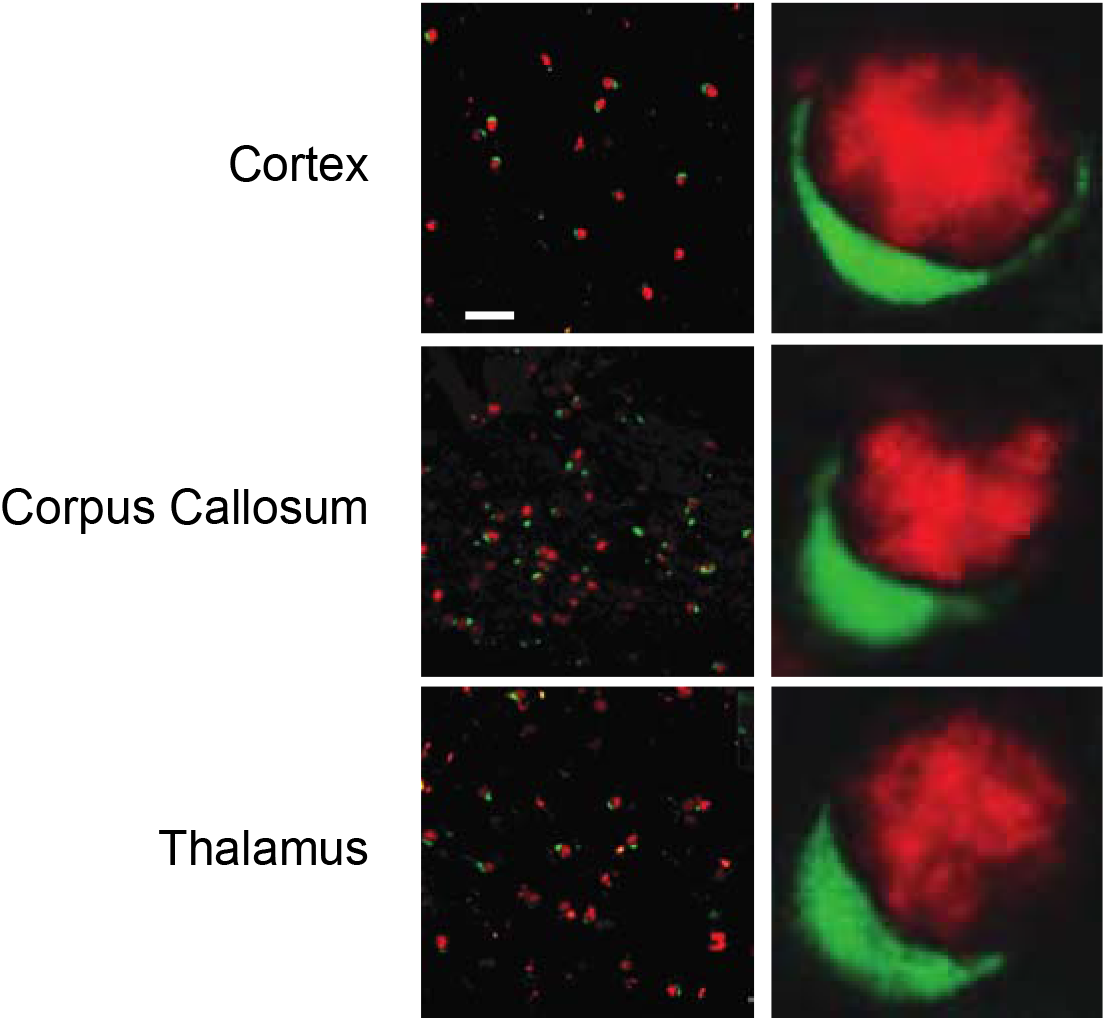
Co-expression of pTau and Olig2 at 180DPI. Parasagittal brain sections from CHI mice were stained for pTau (PHF1, green) and the oligodendrocyte-specific nuclear protein Olig2 (red). Left, representative images of co-labeled PHF1^+^, Olig2^+^ cells in cortex, corpus callosum, and thalamus at 180 DPI. Scale bar, 25μm. Right, high power images of co-localized cells from each brain region.

### Increased thalamic thioflavin-S^+^ cell density at 180 DPI

Thioflavin stains β-pleated sheet structures (Santa-María et al., 2006). Mice expressing a tau mutant P301L allele in oligodendrocytes first develop pTau perinuclear aggregates that later stain with thioflavin-S (Higuchi et al., 2005). Thioflavin-S^+^ cell density was assessed in ipsilesional cortex, corpus callosum and thalamus at 14 and 180 DPI (Figure 6). Thioflavin-S^+^ cell density in cortex or corpus callosum had no significant effect of injury, time or injury*time. In contrast thioflavin-S^+^ cell density in thalamus had a significant effect of injury, time and injury*time. At 180 DPI, thioflavin-S^+^ cell density in thalamus was significantly higher than at 14 DPI (p<0.001). These data suggest an injury, age and region-dependent accumulation of thioflavin-S^+^ cell density. Similar perinuclear structures in ipsilesional thalamus were stained with anti-pTau antibodies (Figures 3, 6A, left inset). Some cells also had thioflavin-S stained processes (Figure 6A, right inset). Ipsilesional thalamic volume correlated highly with thioflavin-S^+^ cell density (r_7_ =-0.90, p=0.005) which was further shown to be a highly linear (F_1,6_=21.8, p=0.005, R^2^ =0.81 (Figure 6C). These data suggest a similar pathological process underlies the increase of thioflavin-S^+^ cell density and thalamic atrophy.

**Figure 6.**
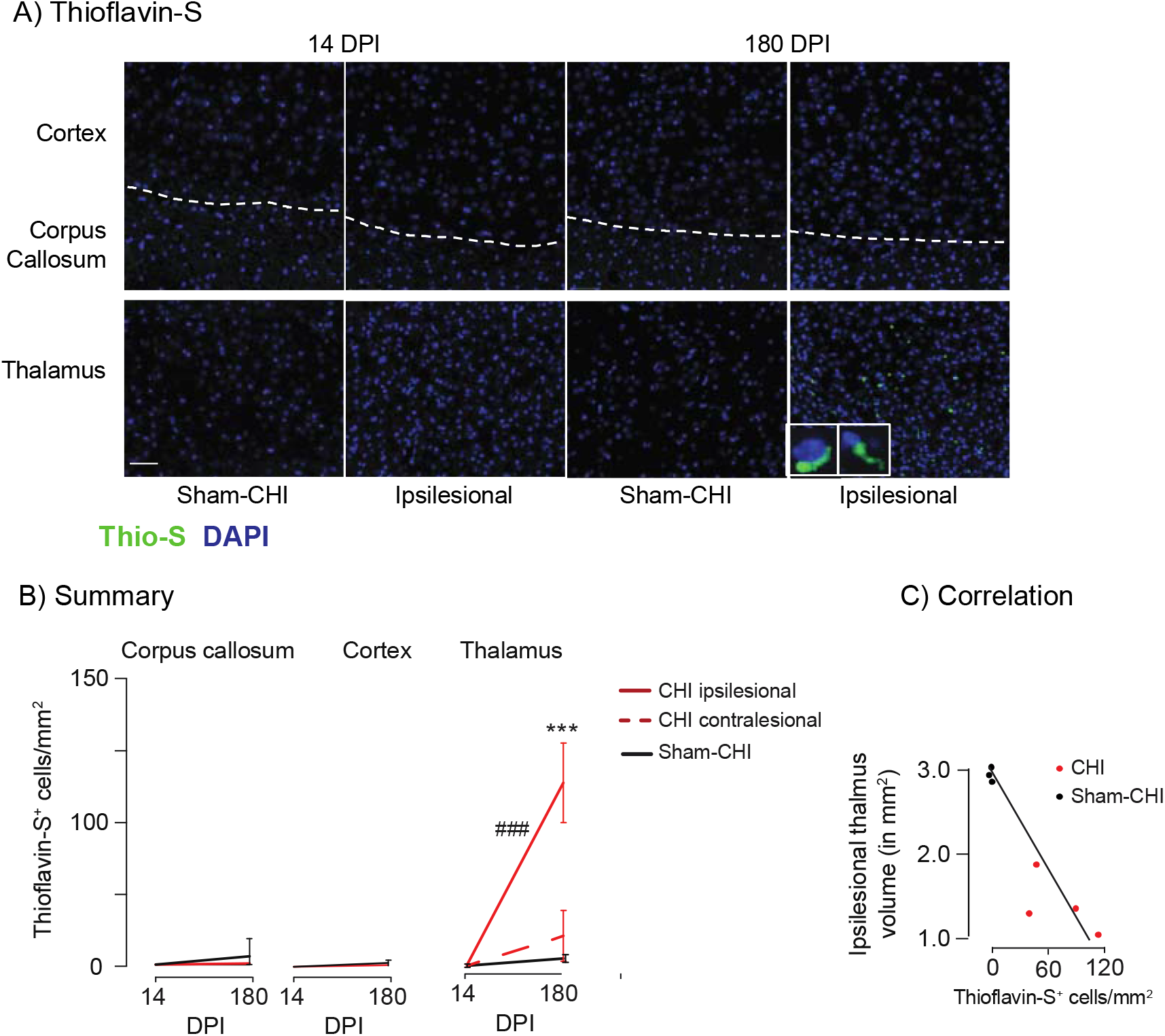
Increased Thioflavin-S^+^ cell density in thalamus. Panel A, Representative images of cortex/corpus callosum (top) and thalamus (bottom) showing staining of thioflavin S-reactive aggregates (thioflavin-S, green) and nuclei (DAPI, blue). Insets if ipsilesional thalamus at 180 DPI show a cell with a thioflavin-S^+^ process (left) or perinuclear aggregate (right). Scale Bar, 50µm. Panel B, Summary of changes in thioflavin-S+ cell density. ***p < 0.001, significant change from sham CHI at 180 DPI, ^###^p<0.001, significant change from CHI at 14 DPI. Panel C, Ipsilesional thalamic volume changes in a linear manner with thioflavin-S^+^ cell density (F_1,6_=21.8, p=0.005, R^2^=0.81).

## Discussion

Acute deficits after a moderate-to-severe TBI may improve with time and rehabilitation, yet many deficits do not improve (Corrigan and Hammond, 2013). A subset of these patients subsequently develop chronic progressive functional deficits (Corrigan and Hammond, 2013). It is poorly understood why some patients improve, while others do not. TBI can produce a chronic neurodegeneration characterized by atrophy of grey and white matter (Bigler, 2021; Cole et al., 2018; Johnson et al., 2017; Wilson et al., 2017). Degeneration and atrophy are likely to be a major driver of behavioral and cognitive decline (Graham and Sharp, 2019).

CHI produces a rapid axonal injury and a chronic injury to gray and white matter (Figures 1, 2). Gray matter atrophy proximal to the impact site occurred within 14 DPI in cortex and thalamus (Figure 2C). This gray matter atrophy was unchanged at 180 DPI (Figure 2C). This is similar to the gray matter atrophy reported in the murine controlled cortical impact (CCI) TBI model that produces rapid gray matter atrophy that continued to increase between 14 and 30 DPI and persists at 180 DPI (Mohamed et al., 2021; Soni et al., 2020).

In clinical TBI, atrophy of white matter is more pronounced than atrophy of gray matter (Cole et al., 2018; Graham and Sharp, 2019). White matter atrophy is more pronounced in the CHI model as well. CHI rapidly induced widespread axonal injury followed by a diffuse white matter atrophy (Figure 1, 2). White matter tracts close to the impact site, such as the ipsilesional corpus callosum and dorsal fornix, show rapid atrophy. More distal white matter tracts, such as contralesional corpus callosum and the bilateral internal capsule and ipsilesional ventral fornix, have delayed atrophy (Figure 2). White matter regions undergoing rapid atrophy likely have white matter microstructural disruptions for they have strongly decreased FA and increased MD and RD (Figures 2B, 2C). In contrast, modest to no change in DTI parameters were seen in white matter tracts showing delayed white matter atrophy (Figure 2B, 2D). Future studies are needed to understand how delayed white matter atrophy occurs with minimal alterations of diffusion tensor parameters.

Injury increased pTau^+^ cell density in white and gray matter (Figures 3,4,7). pTau^+^ cells were exclusively in the oligodendrocyte lineage (Figure 5). This expression pattern of pTau is consistent with limited clinical studies reporting that a single moderate-to-severe TBI induces tau accumulation in oligodendrocytes (Irving et al., 1996; Smith et al., 2003). Accumulation of pTau in oligodendrocytes clearly differs from CTE in which mild repetitive TBIs induce neuronal and astrocytic pTau accumulation (McKee et al., 2016; McKee et al., 2013). The presence of oligodendrocytic pTau rules out CTE as a single diagnosis, as determined by a consensus panel from the National Institutes of Neurological Disorders and Stroke (McKee et al., 2016). This suggests that the tauopathy produced by a single TBI may share pathophysiology with other tauopathies that accumulate pTau accumulation in oligodendrocytes including corticobasal degeneration, progressive supranuclear palsy, and frontotemporal dementia with parkinsonism-17 (Kahlson and Colodner, 2015; Richter-Landsberg, 2016).

Adult human neurons express six tau protein isoforms; that contain either 3 or 4 microtubule binding domains (3R or 4R) (Wang and Mandelkow, 2016). Adult mouse oligodendrocytes express both 3R and 4R tau; but only express 4R tau in neurons (Lopresti, 2002). Differences in tau isoform expression between mice and humans may explain the absence of pTau-expressing neurons following CHI (Lopresti, 2002). Alternatively, CHI may induce pTau aggregation on both neurons and oligodendrocytes prior to 14 DPI followed by selective loss of pTau expressing neurons. This possibility is supported by the finding that CHI induces neuronal loss by 14 DPI (Whitney et al., 2021). Similarly, experimental stroke induces rapid neuronal loss with a longer-lasting pTau deposition in oligodendrocytes (Valeriani et al., 2000).

Phosphorylation at the sites recognized by AT8, PHF1 and S214 disrupts the binding of tau to microtubules. (Goedert et al., 1995; Illenberger et al., 1998). Tau stabilizes the microtubules in the myelinating projections of oligodendrocytes (LoPresti, 2018; Richter-Landsberg, 2008). Decreased tau binding may destabilize and depolymerize microtubules in the cytoskeleton of oligodendrocytes (LoPresti, 2015). Oligodendrocytes form a neuron-oligodendrocyte unit with myelinated axons; disruption of this unit can lead to CNS degeneration (LoPresti, 2018; Richter-Landsberg, 2016). CHI-induced axonal injury likely disrupts the neuron-oligodendrocyte unit, causing the withdrawal of myelinating oligodendrocytic projections (Figure 1)(Richter-Landsberg, 2016). Thus, oligodendrocytic pTau accumulation may lead to cytoskeletal disruption and white matter atrophy (Figures 2 and 3) (Richter-Landsberg, 2016).

At 180 DPI, Thioflavin-S stained perinuclear aggregates in the ipsilesional thalamus, but not cortex or corpus callosum (Figure 6). pTau antibodies stained similar aggregates in thalamus at both 14 and 180 DPI, suggesting formation of pTau-containing perinuclear aggregates (Figures 3, 6). Thioflavin-S^+^ cell density significantly correlated with thalamic atrophy suggesting a common pathological process. Interestingly, axonal injury, thalamic atrophy, and pTau deposition preceded the appearance of Thioflavin-S reactive aggregates (Figures 1,2,3).

Atrophy of ipsilesional corpus callosum is accompanied by an increase in total cell density. This suggests that decreased white matter volume underlies increased cell density. Atrophy without changing cell number in ipsilesional corpus callosum likely contribute to the altered diffusion parameters as measured by DTI (Figure 2).

This study reveals the complexity of chronic neurodegeneration after a single CHI. In the ipsilesional hemisphere, atrophy in the cortex, corpus callosum, thalamus and dorsal fornix occurred within 14 DPI. Atrophy in the internal capsule, and ventral fornix was delayed and progressive. Between 14 and 180 DPI, pTau^+^ cell density decreased in cortex and thalamus, but increased in corpus callosum. Thioflavin-S^+^ cell density increased only in thalamus. In the contralesional hemisphere, atrophy in the cortex, corpus callosum, and internal capsule was delayed and progressive. In the ipsilesional hemisphere, pTau^+^ cell density never increased in cortex, but increased corpus callosum (Figure 3). Thus, chronic neurodegeneration in individual brain regions differs depending on their proximity to the impact site and whether they are gray or white matter.

There are important caveats to this study. No assessments were made between 14 and 180 DPI; a more detailed time course will likely increase our understanding of how atrophy, pTau deposition and thioflavin-S^+^ cells progress over time. Similarly, no studies have been done past 180 DPI to determine if these neurodegenerative changes are ongoing. This study used only used 5 antibodies against pTau epitopes, which provides limited insight into the pathogenic potential of tau. Chronic inflammation is well established in both clinical TBI and TBI models; a better understanding of inflammation in the CHI model could provide further information regarding why CHI can produce multiple forms of chronic neurodegeneration (Jassam et al., 2017). Despite these caveats, this study clearly shows that a single CHI produces a profound chronic, progressive neurodegenerative pathology in both gray and white matter.

## Acknowledgements

We thank members of the Neuropathology Brain Bank & Research CoRE at the Icahn School of Medicine at Mount Sinai (New York) for histological assistance.

---

**Supplementary Figure 1.**
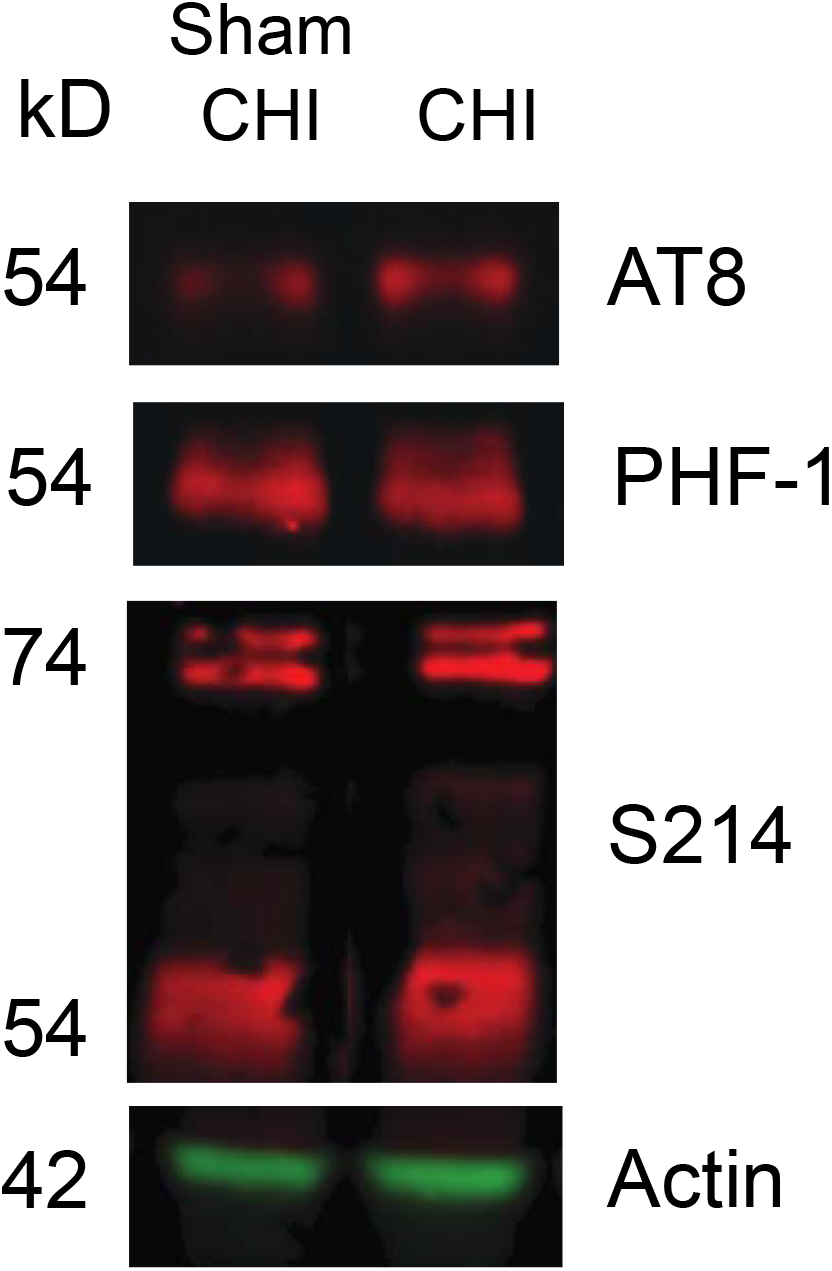
pTau protein levels in thalamus are unchanged by CHI. Protein was extracted from the thalamus of the sham-CHI and CHI groups at 14 DPI and immunoblotted against AT8, PHF1 and S214 by the method of Rubenstein, et al., (Rubenstein, 2015). AT8 reacted with a 54kD protein species in both the sham-CHI and CHI groups. The amount of 54kD protein was similar in both the sham-CHI and CHI group (t_5_ =0.64, p = 0.55). PHF1 reacted with 55.5 and 55.1 54kD protein species. The amount of the 55.5 KD immunoreactive band was 28.1 ± 13.1% lower in the CHI group than the sham-CHI group (t_6_ = 2.63, p = 0.03). In contrast, the amount of the 55.1 kD immunoreactive band was similar in the CHI and sham-CHI groups (104.28 ± 10.45%, t_6_ = 0.39, p = 0.71). S214 reacted with 77.1, 73.8, 53.3, and 51.7 kD immunoreactive bands. A similar pattern of S214 immunoreactive proteins of 70-80 kD have been reported by others (Wang, et al., 2018 Zhou, et al., 2018). The amounts of 77.1, 73.8, 53.3, and 51.7 kD S214 immunoreactive bands were similar in the CHI and sham-CHI groups (78.8 kD, 110.08% ± 15.52%, t_6_ = 1.06, p=0.33; 77.1 kD, 78.34 ± 29.73%, t_6_ = 1.56, p=0.17; 53.5 kD, 83.34 ± 10.20% t_6_ = 1.06, p=0.33; 51.7 kD, 114.85 ± 23.55%; t_6_ = 1.56, p=0.8). These data suggest that CHI did not change in the amount of pTau protein in the thalamus at 14 DPI.

